# Resistance to Predation is Not Constrained by Growth-Defense Trade-Offs in Freshwater Bacterial Communities

**DOI:** 10.64898/2026.11.26.769432

**Authors:** Nikesh Dahal, Vincent J. Denef

## Abstract

Predation is a fundamental force shaping ecological communities through shifts in abundance and trait distribution patterns. Classical theory predicts that investment in predation defense incurs growth costs due to allocation trade-offs. However, whether microbial communities in nature exhibit such trade-offs remains unclear. We test this prediction in bacterial communities of Lake Erie, a high nutrient system under strong top-down control by Dreissenid mussels: ecosystem engineers altering global freshwater. We find that mussel predation drives functional trait shifts, with distinct patterns of both convergence and divergence emerging at phylogenetic, metabolic, and molecular levels. However, we find no evidence of a growth-defense trade-off. Instead, out statistical modelling results suggest that environmental heterogeneity may relax trade-off constraints, favoring resistance strategies that extend beyond classical life-history predictions. Our findings provide evidence that microbial resistance is a complex trait structured across multiple levels of biological organization, with implications for food web ecology and eco-evolutionary theory.

## Introduction

Animal-microbe interactions play a key role in shaping eco-evolutionary processes, affecting physiological responses such as the timing of life-history events (e.g., metamorphosis and reproduction) (Knutie *et al*. 2017; Metcalf *et al*. 2019) as well as fitness outcomes through local adaptations involving life-history trade-offs (Gould *et al*. 2018; Walters *et al*. 2020). While there has been a growing interest in this topic in recent decades, most research has focused on host-microbiome systems, examining the effects of microbiome on animals (McFall-Ngai *et al*. 2013; Mazorra-Alonso *et al*. 2021). In contrast, studies on how interactions involving animals in non-host contexts shape microbial community structure and trait distributions are relatively underexplored, despite their potential to reveal key insights about biodiversity maintenance.

Recent evidence from zoogeochemistry, a field studying the role of animals in biogeochemical cycles, show that animals modulate microbial processes across ecosystems, including in the world’s largest freshwater lakes (Vanni 2002; Schmitz *et al*. 2018; Li *et al*. 2021). These findings challenge the long-held assumption about the primacy of microbial interactions in driving energy flow and nutrient cycling (Falkowski *et al*. 2008), and instead suggest that animal-microbe interactions in non-host contexts can mediate ecosystem dynamics by restructuring microbial functional traits (Schmitz & Leroux 2020). This mediation may occur indirectly, or directly through top-down effects of animal interactions. Top-down effects of animals, thus, represent a fundamental process structuring ecological communities more broadly, as they regulate population densities (Hairston *et al*. 1960; Paine 1966) and trait distributions of the interacting species (Oksanen *et al*. 1981; Werner 1986).

A limited number of studies have explored top-down effects of animals on microbes (Crowther *et al*. 2013; Song *et al*. 2023; Tobias-Hünefeldt *et al*. 2021). Moreover, the strength and direction of these effects can vary across space and time (Denef *et al*. 2017; Dahal *et al*. 2023), underscoring the importance of environmental heterogeneity in shaping microbial response dynamics. Even over short timescales, such variations have been linked to shifts in microbial community functions, such as respiration and nutrient cycling (Braeckman *et al*. 2010; Wyness *et al*. 2021). This suggests that animal-driven selection pressures on microbial functional traits are not uniform but rather emerge from dynamic eco-evolutionary processes. Yet, how this top-down force shapes microbial functional diversity patterns remains poorly understood. It is particularly important to resolve this today, as trophic downgrading (Estes *et al*. 2011) and species introductions (Havel *et al*. 2015) continue to form novel community structures (Vandermeer & Perfecto 2023) with complex and dynamic roles of animal-microbe interactions.

### Trait-Based Approaches in Microbial Community Ecology

In this context, trait-based approaches have emerged as one of the primary tools in ecology. By focusing on shifts in the distribution of functional traits, these frameworks extend beyond pairwise interactions and offer the promise of abstracting generalizable patterns to link community variation with ecosystem processes (Grime 1977; McGill *et al*. 2006). Among these, life-history trade-offs have become a dominant approach in microbial ecology (Lajoie & Kembel 2019; Malik *et al*. 2020). For example, these frameworks have been used to predict phytoplankton community diversity (Litchman & Klausmeier 2008), ecosystem processes like decomposition (Allison 2012), and microbial responses to environmental stressors such as drought (Scales *et al*. 2022). At its core, life-history theory centers on fitness optimization in resource-limited environments, offering an eco-evolutionary lens to link demographic and adaptive processes under equilibrium-based assumptions. Thus, the theoretical expectation under this premise is that energetic constraints drive the emergence of trade-offs between reproductive and survival traits, leading to co-variation of traits along certain niche axes, such as growth and defense (Grime 1977; Malik *et al*. 2020). Grouping species based on life-history strategies can then be used to gain insights about how variation in community structure translates into distribution of functional traits (Adler *et al*. 2014; Ramin & Allison 2019).

However, these frameworks largely assume that organisms invest in fixed strategies, often overlooking the complexity of species interactions in food web contexts, particularly in microbial systems. For example, predator foraging behavior, prey vulnerability, attack rates, and prey movement complicate the generalizability of trait-based frameworks in food webs involving animals (Gravel *et al*. 2016). Similarly, in microbial communities, metabolic flexibility, horizontal gene transfers, and frequency-dependent interactions shape phenotypic and genotypic diversity in ways that fundamentally differ from ones observed in larger organisms (Rodriguez-Valera *et al*. 2009; Polz *et al*. 2013; Lennon & Denef 2015). Despite the widespread use of trade-off frameworks in microbial ecology, few studies have explicitly tested their assumptions in non-host animal-microbe systems, where the complexity of selection landscapes likely disrupts long-term equilibrium-based predictions. A recent meta-analysis by McCary & Schmitz (2021) demonstrated that while functional traits offered a useful framework for understanding invertebrate-mediated nutrient cycling, high context-dependency contributed to significant uncertainty regarding the role of animal interactions in mediating microbial processes. Part of this uncertainty likely stems from how eco-evolutionary processes in non-host animal-microbe interactions shape functional diversity patterns, highlighting the need for empirical studies to test theoretical predictions in these systems.

### Study system

In this study, we focus on Dreissenid mussels, a well-known introduced ecosystem engineer (Strayer 2009; Higgins & Vander Zanden 2010), to investigate how their predation affects freshwater bacterial trait diversity patterns. Dreissenid mussels exemplify the zoogeochemical influence of introduced species, exerting strong controls on microbial processes in freshwater ecosystems (Vanni 2021). For instance, they regulate phosphorus flux at rates ten times higher than natural sedimentation in the Laurentian Great Lakes (Li *et al*. 2021). Lake Erie, one of the earliest North American ecosystems where Dreissenids established populations (Berkman *et al*. 1998), provides an ideal setting to examine the top-down effects of mussel predation on microbial traits. Several factors make this system well-suited for such an investigation: (*a*) mussels have imposed decades of selective pressures on microbial communities (Arnott & Vanni 1996; Vanderploeg *et al*. 2001) (*b*) their predation has been linked to shifts in the phytoplankton (Tang *et al*. 2014; Carrick *et al*. 2023) and bacterial community composition (Lee *et al*. 2015), including proliferation of toxic cyanobacterial blooms dominated by *Microcystis* (Vanderploeg *et al*. 2001) *(c)* predation-driven changes in bacterial diversity are context-dependent (Dahal *et al*. 2023), and *(d)* mussel densities and distribution have varied over time, with the ecosystem exhibiting strong seasonal fluctuations (Carter *et al*. 2023; Vanderploeg *et al*. 2023). Hence, if the growth-defense trade-off is a robust principle under natural conditions, we may expect mussel predation to structure resistant and susceptible bacterial populations along the growth-defense trade-off axis.

Based on the above insights from the natural history of Lake Erie, we explored the effects of mussel predation on bacterial functional traits. Specifically, we used a set of lake water mesocosm experiments with or without mussels that we previously analyzed at a community composition level (Dahal *et al*. 2023). Here, we used metagenomic and statistical modelling approaches to explore how resistance to mussel predation is structured across distinct levels of functional organization - phylogenetic, metabolic, and protein levels. We hypothesized that observed shifts in bacterial community structure and composition would manifest as growth-defense tradeoffs among resistant and susceptible populations. While we observed distinct patterns of functional shifts at each level of organization, we found no evidence for the existence of a growth-defense trade-off. Instead, our statistical modelling results suggest that environmental heterogeneity in Lake Erie likely relaxes the constraints predicted under the equilibrium conditions.

## Results

We recovered 214 metagenome assembled genomes (MAGs) from Lake Erie bacterial communities after manually curating 298 high quality bins (> 80% completion and <10% redundancy) generated through an automated binning approach. After dereplication of highly similar MAGs, 181 unique MAGs were retained for analyzing resistance to mussel predation as a continuous trait (Figure 1). A subset of 149 MAGs met the criteria for categorical analysis, with 15 resistant and 134 susceptible populations based on relative abundance shifts in response to quagga mussel predation. All MAGs exhibited high genome completeness (mean = 91.08%) and low redundancy (mean = 3.41%), indicating that they reflect well-assembled representations of natural bacterial populations from Lake Erie.

**Figure 1.**
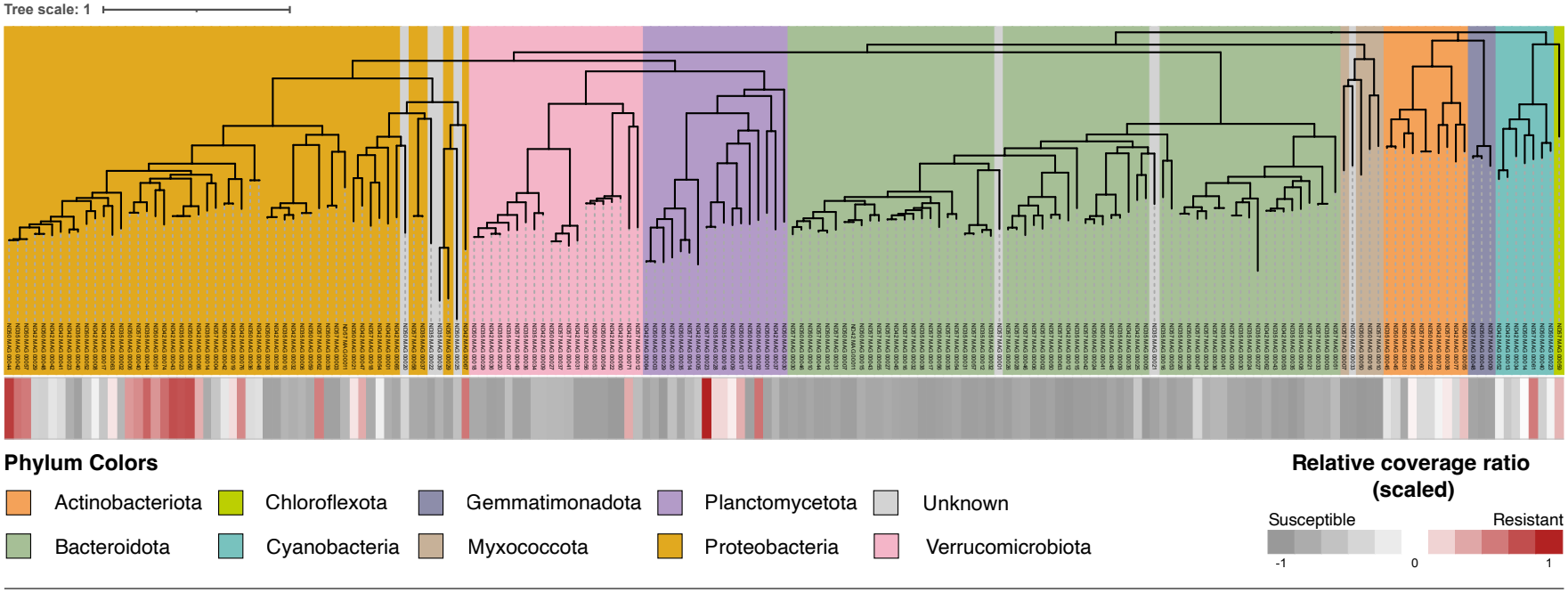
Phylogeny of metagenome assembled genomes. A maximum-likelihood phylogenetic tree of 181 metagenome assembled genomes (MAGs) recovered from Lake Erie bacterial communities. Background shading indicates taxonomic distribution at the phylum level. Heatmap panel (below the tree) displays mean resistance across experimental dates for each MAG, calculated as the relative abundance ratio in the quagga treatment (Q) vs. control treatment (C) and scaled between -1 and 1 for visualization. Mantel test revealed a weak but significant correlation between phylogenetic structure and resistance trait distance (r = 0.0487, p-value = 0.047, 999 permutations).

The average GC content was 53.43% with values ranging from 29.67 - 74.36%, reflecting phylogenetic and functional diversity represented in our dataset. Taxonomically, populations were distributed across the same phyla previously identified in our 16S rRNA gene-based analyses of these experiments - *Proteobacteria, Bacteroidota, Verrucomicrobiota, Planctomycetota, Actinobacteriota, Cyanobacteria, Myxococcota, Gemmatimonadota*, and *Chloroflexota* - confirming consistency in our metagenomic characterization of the ecological communities (Dahal *et al*. 2023). Given that dereplicated MAGs were used to delineate bacterial lineages, *MAGs* and *populations* are used synonymously in the following sections. A full descriptive summary of these populations is provided in Table S1.

### Phylogenetic Patterns

Resistance to mussel predation exhibited a strong phylogenetic signal in the Lake Erie bacterial communities (Pagel’s λ = 0.988), suggesting that the evolution of this trait does not follow a neutral Brownian motion model and closely related populations tend to respond similarly to predation (Figure 1). However, the Mantel test revealed only a weak but significant correlation between phylogenetic structure and trait distance (r = 0.049, p-value = 0.047, 999 permutations), indicating that genetic divergence alone does not explain the observed phylogenetic clustering. A phylogenetic generalized least squares (PGLS) model accounting for evolutionary history revealed that GC content was a significant predictor of resistance, although it explained minimal variation (β = -0.010, p-value = 0.011), whereas genome size had no significant effect (p-value = 0.327). Phylogenetic signal remained strong (λ = 0.987) in this model, reinforcing the role of evolutionary constraints in shaping the trait distribution. Consistent with the PGLS results, Wilcoxon rank sum tests conducted for each sampling date between resistant and susceptible populations did not reveal significant differences in GC content or genome size, (all p-value > 0.05).

### Functional differences between resistant and susceptible populations

Among the 4,226 shared protein families (pfams), 23 showed significant differences in their mean relative abundance between resistant and susceptible populations (BH-adjusted p-value < 0.05; Figure S1A). Of these, 21 pfams were overrepresented in resistant populations and 2 were more abundant in susceptible populations (Figure 2A). Key pfams with higher differential abundance in resistant populations include ABC transporters (PF00005.31), binding-protein-dependent transport systems (PF00528.26), toxin-antitoxin systems (PF13304.10), and oxidative stress response proteins (PF13450.10, PF07992.18) (Figure 2B). Principal Coordinates Analysis (PCoA, Bray-Curtis) of these 23 pfams did not reveal discrete clustering between resistant and susceptible populations (Figure 2C). There was a sharp drop in eigenvalue after PC1 (Figure S1B), with 60% of variance represented on the first axis. PERMANOVA analysis detected a significant taxonomic signal (p-value = 0.002; Figure S1C), suggesting that pfam composition varies by phylum for the resistant populations. However, this result was confounded by uneven taxonomic structure, as most resistant populations belonged to the *Proteobacteria* phylum, and significant dispersion differences (ANOVA, p-value = 0.012) indicate that variation within phyla also contributed to the observed separation.

**Figure 2.**
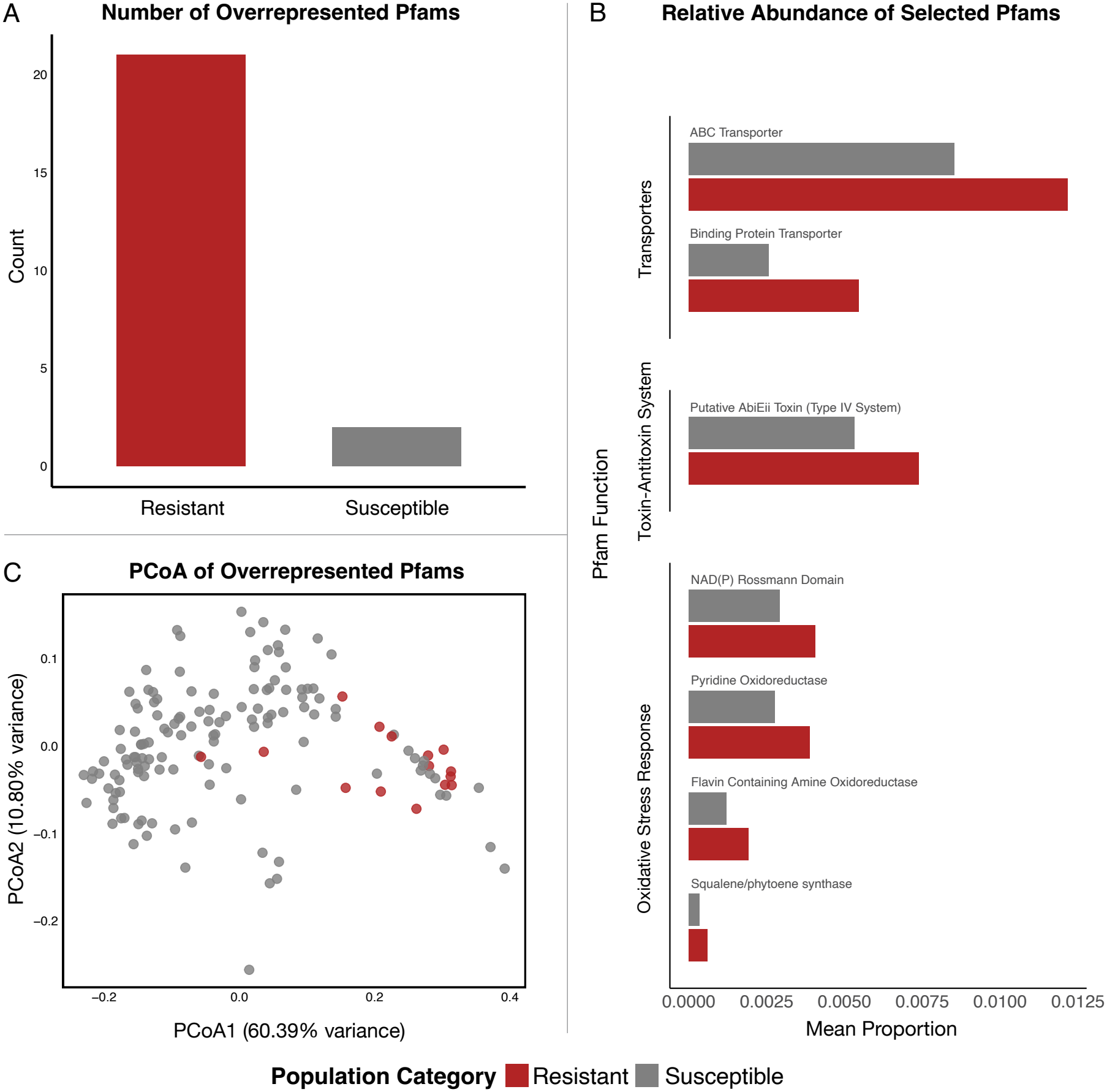
Overrepresented protein families and their composition. (A) Differential abundance of protein families (pfams) between resistant and susceptible populations, with 21 out of 23 functions overrepresented in resistant populations (B) Relative abundance of key pfams associated with resistance to mussel predation categorized into three functional groups: transporters (top panel), toxin-antitoxin systems (middle panel), and oxidative stress response proteins (bottom panel). Bars represent mean relative abundance in resistant (red) and susceptible (gray) populations, highlighting functional diversity in protein level resistance mechanisms. (C) Principal Coordinates Analysis (PCoA, using Bray-Curtis) of both populations based on the 23 enriched pfams. Resistant and susceptible populations did not form distinct clusters, indicating continuous rather than discrete variation in pfam composition associated with resistance to mussel predation.

Of the 35 KEGG metabolic modules that differed significantly between resistant and susceptible populations, 26 were more abundant in resistant populations. These included functions related to amino acid metabolism, energy metabolism, and nucleotide metabolism (Figure S2A). Resistant populations also had a significantly higher metabolic module completeness (Wilcoxon rank sum test, p-value < 0.001; Figure 3A). Parallel to KEGG, comparative analyses of clusters of orthologous groups (COGs) also revealed functional differentiation, with five out of eight COG categories overrepresented in resistant populations (BH-adjusted p < 0.05; Figure S2B), including amino acid transport and metabolism, energy production and conversion, and mobilome-associated elements. Notably, amino acid and energy metabolism were consistently overrepresented in resistant populations across both, KEGG and COG, datasets (Figure 3B). To assess whether these functional traits are structured by taxonomy, we conducted PCoA (Bray-Curtis) of overrepresented KEGG modules within energy and amino acid metabolism - key functions overrepresented in resistant populations. Like the earlier pfam analysis, PCoA revealed marginally significant clustering by taxonomic groups (PERMANOVA, p-value = 0.062; Figure S3C) but this separation was confounded by taxonomic dispersion (ANOVA, p-value = 0.043).

**Figure 3.**
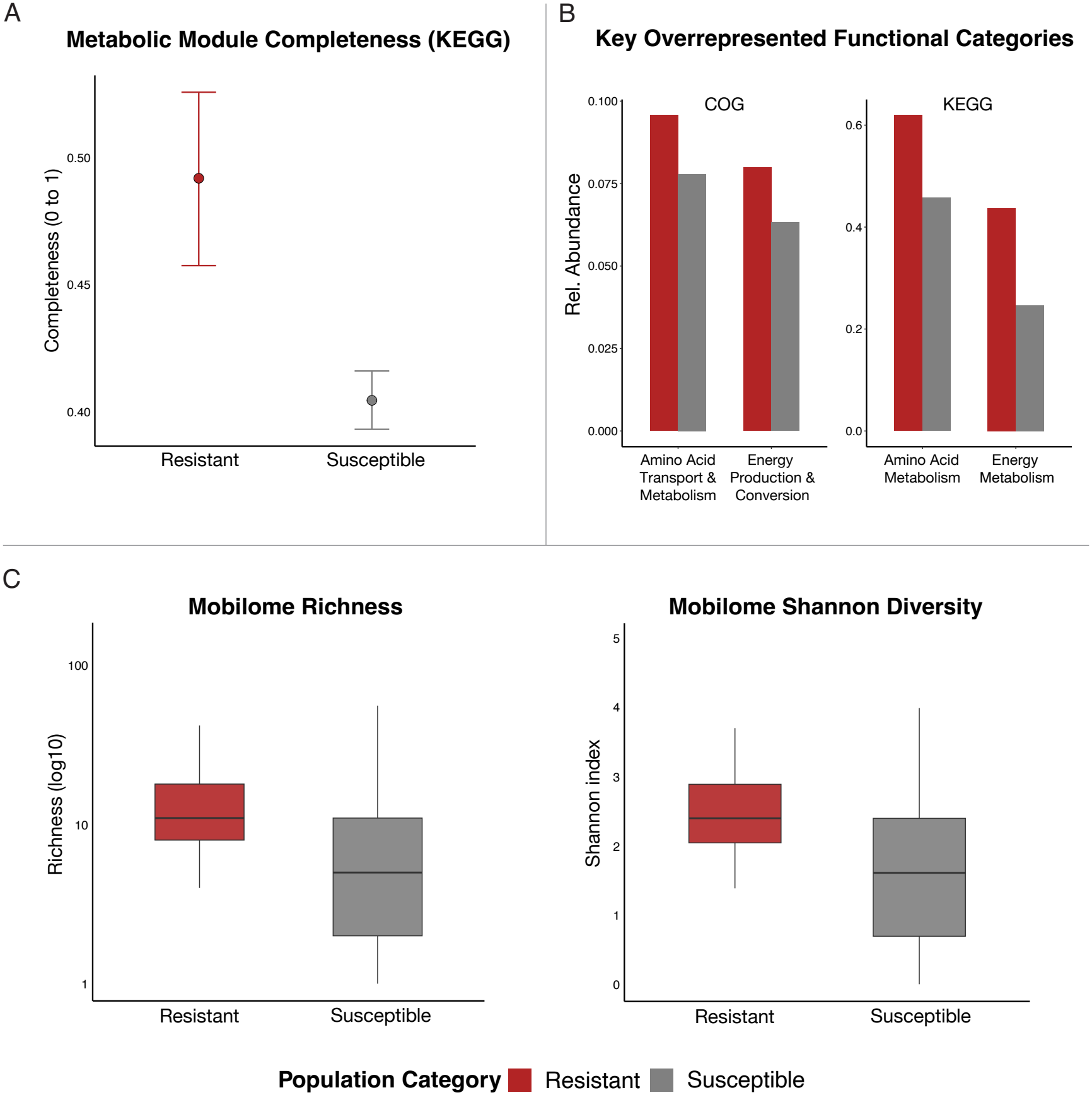
Differences in metabolic pathway completion, broad functions, and mobile genetic elements. (A) Metabolic module completeness of resistant (red) and susceptible (gray) populations. Mean completeness (±95% CI) is higher in resistant populations (p < 0.001, Wilcoxon test), indicating functional differentiation in the metabolic niche space. (B) Relative abundance of COG categories (left) and KEGG modules (right) for amino acid and energy metabolism. Both annotation approaches showed overrepresentations of these metabolic pathways in resistant populations, indicating overlap in the metabolic niche space at the level of core metabolic functions and conserved functional genes. (C) Comparison of the mobilome diversity between two populations. Resistant populations have significantly higher mobilome richness (p-value = 0.008) and Shannon diversity (p = 0.008; Wilcoxon rank sum test), indicating greater variability in mobile genetic elements such as transposons and prophases.

While pfam, KEGG, and COG analyses did not indicate enrichment of secondary metabolite production, antiSMASH identified biosynthetic gene clusters (BGCs) in resistant populations. We detected 26 BGCs across resistant populations (Table S4), encoding terpenes, Type III polyketide synthases (PKS), and nonribosomal peptide synthases (NRPS), which are chemical classes known to be involved in microbial defense mechanisms (Rohlfs & Churchill 2011).

### Mobilome diversity and metabolic capacity

Beyond core metabolic differentiation, we found that resistant populations had a significantly higher diversity in their mobilome - a suite of mobile genetic elements, including transposons and prophages, which move within and between genomes and facilitate horizontal gene transfer (Wilcoxon rank sum test: richness, p value = 0.007; Shannon diversity, p value = 0.008; Figure 3C). Moreover, mobilome diversity was positively correlated with functional capacity, indicating that populations with higher mobilome Shannon diversity index tend to have a greater number of unique metabolic modules (Spearman’s ρ = 0.36, p < 0.001; Figure S3).

### Growth-defense trade-offs and population persistence

We found no significant difference in maximum growth rate estimates of populations resistant and susceptible (using gRodon) to mussel predation (Wilcoxon rank sum test, p-value = 0.302) (Figure 4A). Bootstrapping for resistant populations and downsampling for susceptible populations (both 1,000 iterations) resulted in overlapping 95% confidence intervals (Figure 4A), suggesting that any observed differences are likely due to stochastic effects rather than adaptive divergence in intrinsic growth-related traits. However, contrary to theoretical expectations, we found that the average instantaneous growth rate was higher for resistant populations compared their susceptible counterparts across 30 randomly selected populations (median irep values = 2.73 vs 2.48; Wilcoxon rank sum test, p-value < 0.001; Figure 4B). When fitting the data to various linear mixed effects models, model 3, which included random slopes for date and treatment and included an interaction term (treatment X population category), provided the best fit to the data (ΔAIC = -253.4, p < 0.001). This model captured a significantly greater proportion of variance in instantaneous growth rates (~90%) and substantially reduced the residual variance compared to model 2 (57%) and model 1 (52%). Moreover, it consistently outperformed model 2 (R^2^: 0.61 to 0.69 vs 0.36 to 0.44) across a range of training dataset (80% to 40% of observed data; Table S3). In short, we find no evidence for the support of growth-defense trade-off predicted under the constraints imposed by fixed investment strategies.

**Figure 4.**
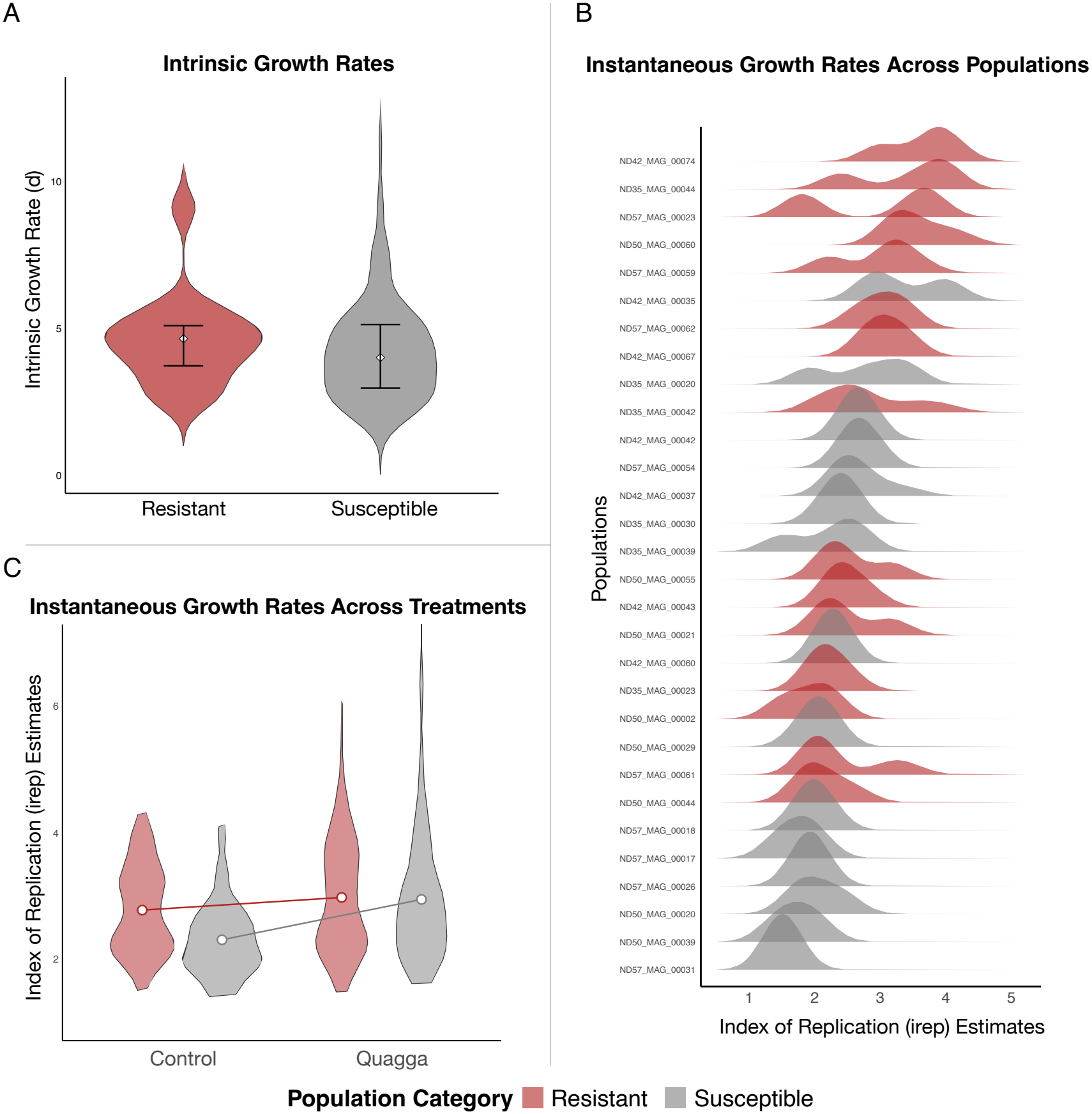
Growth rate comparisons between resistant (red) and susceptible (gray) populations. (A) No significant difference in intrinsic growth rates (Wilcoxon rank sum test, p-value = 0.2246). (B) Growth rate distributions varied substantially between the two populations when pooled across dates and experimental treatments. (C) Resistant populations exhibited significantly higher instantaneous growth rates (Wilcoxon rank sum test, p-value < 0.001), contradicting expectations based on growth-defense trade-off theory. Statistical modelling revealed a significant interaction between the experimental treatment and population category (p < 0.001; Table S2).

Lastly, building on the role of temporal (date) effects, we examined how resistance was associated with population persistence. A chi-square test revealed significant association between population persistence and resistance (X2 = 6.07, df = 2, p-value = 0.048). Populations persisting across all three dates had the highest proportion of resistant lineages (~29 %) compared to those observed in two dates (~20 %) or only one (~9 %) (Figure 5). Notably, the number of bacterial populations in all three categories was similar (N = 40-54), suggesting that the observed association between persistence and resistance likely reflects natural population and community dynamics rather than sampling bias.

**Figure 5.**
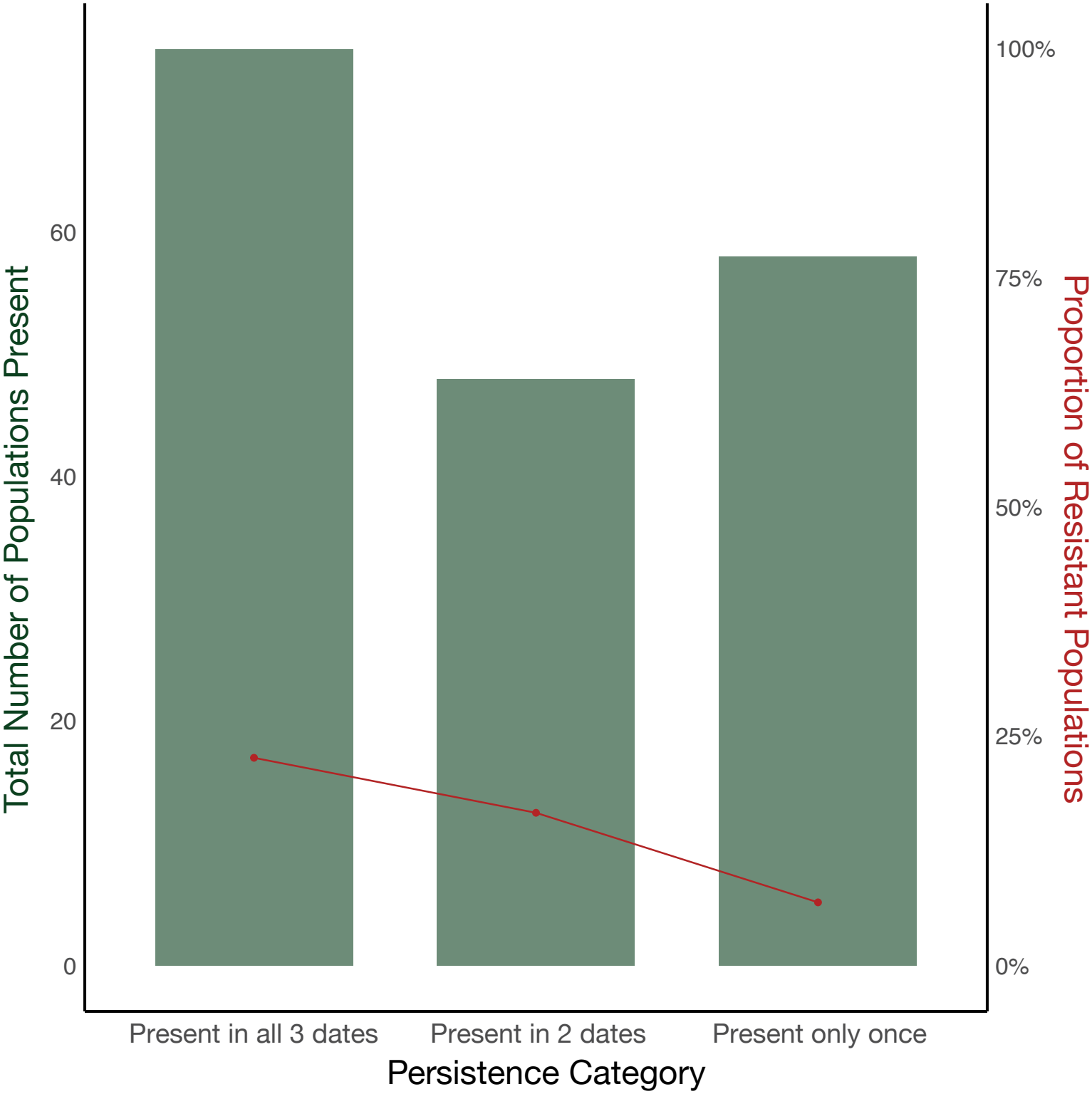
Proportion of resistant populations: Total number of populations (green bar, left Y-axis) and proportion of resistant populations (red line, right Y-axis) grouped by persistence across sampling dates (observed in all three dates, two dates, and once). A significant association between population persistence and resistance (chi-square, X2 = 6.07, df = 2, p-value = 0.048), suggests that resistant populations are more frequently observed across time compared to susceptible populations.

## Discussion

Classical life-history theory predicts that investment in defense traits comes at the cost of growth due to energetic constraints (Stearns 1989; Tilman 2000). However, our results contradict this prediction: (a) intrinsic growth rates did not differ between resistant and susceptible bacterial populations (Figure 4A), and (b) resistant populations exhibited higher instantaneous growth rates, counter to the predictions (Figure 4C). Given that Dreissenid mussels have imposed strong ecological selection on Lake Erie microbial communities for decades (Vanderploeg *et al*. 2001, 2009), we expected resistance to mussel predation to be constrained by growth-defense trade-offs. Instead, we saw that this trait was structured across multiple scales of functional organization, with different patterns emerging at phylogenetic (Figure 1), metabolic (Figure 3), and molecular (Figure 2) levels. Moreover, resistant populations were generally more persistent than more abundant, compared to susceptible populations (Figure 5)(Dahal *et al*. 2023), and overall exhibited greater metabolic independence (Figure 3A). This suggests that resistance traits are associated with survival in heterogeneous environments rather than local competitive dominance, as organisms adapt to patterns and processes of their environment rather than fitting into static niches (Levins 1979). This aligns with recent findings in microbial ecology, where rare but persistent populations with high metabolic independence have been identified as an emergent property of microbial communities in stressful environments (Watson *et al*. 2023; Veseli *et al*. 2024).

### Why did we not observe patterns supporting the growth-defense trade-off framework?

Equilibrium-based abstractions underpin growth-defense trade-off theory, yet theoretical and empirical studies reveal that ecological communities rarely exist in equilibrium (Allesina & Levine 2011; Huisman *et al*. 2018). In fact, it has been shown that non-equilibrium dynamics involving species interactions, including predation, hold immense potential in generating and maintaining biodiversity patterns (Scheffer *et al*. 2003; Vandermeer & Perfecto 2024), which may help explain why trade-offs did not structure observed diversity patterns in our study. Our study system, Lake Erie, is a nutrient rich lake where seasonal variation in temperature, nutrient availability, toxic cyanobacterial blooms, and phytoplankton succession create a spatiotemporally heterogeneous matrix (Vanderploeg *et al*. 2023), likely disrupting equilibrium-based constraints. Consistent with this, our statistical modelling results (Table 1, S2, S3) and experimental data (Figure 4) indicate that temporal heterogeneity is a crucial factor shaping bacterial growth responses to mussel predation. Hutchinson’s Paradox of the Plankton (1961) offers a useful conceptual parallel in this context: if microbial communities adapt to variability in the environment rather than optimizing fixed investments along resource allocation trade-offs, then dynamic metabolic strategies can emerge and maintain diversity rather than constrain it through competitive exclusion (Pacciani-Mori *et al*. 2020).

**Table 1.**
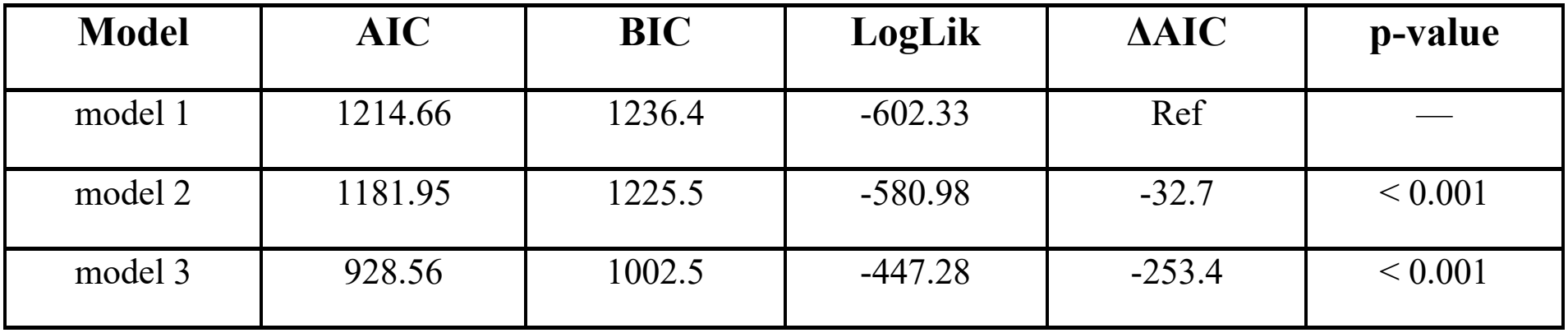
Comparison of three linear mixed-effects models used to fit the observed instantaneous growth rate estimates from iRep. Models are ranked by Akaike Information Criterion (AIC), with lower values indicating better model fit. Bayesian Information Criterion (BIC) and log-likelihood (LogLik) values are also reported. ΔAIC represents the difference in AIC relative to the baseline model (model 1). Model 3, which includes random slopes for treatment and date, along with an interaction term for the experimental treatment and population category, provides the best fit (ΔAIC = -253.4, p < 0.001; Table S2), significantly outperforming model 1 and model 2 (Table S3).

Resistance to mussel predation lacked significant correlation with genome length, GC content and intrinsic growth rate (Figure 4A), and exhibited only a weak phylogenetic signal indicating that it is not a deeply structured evolutionary trait. Nonetheless, the distribution of resistant populations across the phylogeny was non-random, exhibiting some level of clustering (Figure 1; Mantel test, r = 0.0487, p-value = 0.047). This suggests that resistance to mussel predation is a complex trait structured beyond single genes but more flexible than highly conserved traits (e.g: salinity preference or photosynthesis)(Martiny *et al*. 2013). Indeed, at the molecular level, resistance was associated with increase in relative abundance of protein families (pfams) involved in oxidative stress response, transporter proteins, and toxin-antitoxin systems (Figure 2A, 2B). Yet, despite this functional differentiation, no differentially abundant pfam was consistently associated with resistance across resistant populations, as PCoA of the overrepresented pfams did not result in a clear separation between the two population categories (Figure 2C). These results, along with the identification of 26 biosynthetic gene clusters—genetic machineries that produce secondary metabolites important for microbial defense (Xia *et al*. 2022)—across resistant populations, suggests that multiple molecular mechanisms underpin this trait (Table S4). While we initially hypothesized that the differences in mechanisms may map onto the phenotypic clusters within resistant populations, the phylogenetic signal was confounded due to taxonomic overdispersion (Figure S1C, Figure S2C). Taken together, our results indicate that interactions between bacterial communities and mussels in Lake Erie are mediated through both evolutionary constraints as well as independently acquired mechanisms, resulting in diverse functional strategies suited to withstand the top-down effects under variable environmental conditions.

Our KEGG and COG analyses similarly revealed an overlap in the metabolic niche space among resistant populations, with amino acid metabolism and energy metabolism emerging as key functional groups associated with resistance among resistant approaches (Figure 3B). While KEGG captures the molecular wiring by mapping modules organizing metabolic pathways (Kanehisa 2019), COG focuses on functional evolution by tracing the genes shared through ancestry (Koonin 2005). These complementary approaches reflect distinct aspects of the hierarchical organization of biological function and highlight convergent metabolic adaptations in resistant populations at a broad scale, despite divergence at a finer scale, as pointed out in the pfam analyses. Additionally, resistant populations exhibited a higher metabolic module completeness (Figure 3A), suggesting that resistance is not only associated with specific molecular functions, but also linked to broader metabolic strategies related to population persistence across changing environmental conditions (Figure 5). These patterns of functional overlap are supported by our previous findings from oligotrophic Lake Michigan. While there are notable differences in the impacts of mussel predation on the microbial diversity patterns in these two systems (Denef *et al*. 2017; Dahal *et al*. 2023), one abundant resistant population that has been genomically examined, Chloroflexi CL-500, exhibited increased expression of oxidative stress response, and amino acid metabolism (Denef *et al*. 2016), matching the metabolic patterns observed in Lake Erie. Thus, we conclude that energy and amino acid metabolism represent key features of the metabolic niche space through which bacterial populations adapt to mussel predation, potentially constraining how resistance emerges within broad metabolic strategies across ecosystems.

The overrepresentation of mobilome associated genes (Figure S2B) in resistant populations, combined with their significantly higher mobilome diversity (Figure 3C), suggests that genome mobility plays an important role in shaping resistance beyond phylogenetic constraints (Acman *et al*. 2022; Botelho *et al*. 2023). Mobilome diversity positively correlates with KEGG metabolic module richness (Spearman’s ρ = X, p < 0.001; Figure S3), reinforcing the idea that genomic rearrangements - whether through horizontal gene transfer (HGT) or within-genome recombination - likely expand the metabolic niche space in resistant populations (Wiedenbeck & Cohan 2011; Szappanos *et al*. 2016). While HGT is well-known to facilitate exchange of adaptive traits across distantly related lineages (Soucy *et al*. 2015), the observed patterns may very well reflect historical acquisitions and loss of the genetic material. This distinction is crucial, as previous studies have indicated that within-genome recombination and habitat structuring contribute to differentiation and coexistence of distinct subpopulations within genotypic clusters (Denef *et al*. 2010; Polz *et al*. 2013). Moreover, previous data on the spatiotemporal distribution of phylogenetically cohesive populations of *Microcystis* in Lake Erie (Berry *et al*. 2017), a resistant population observed in our experiments, suggests that fluctuating ecological conditions are likely important in shaping the observed genomic patterns.

While our study provides key insights into how resistance to mussel predation shapes bacteria community diversity, we acknowledge its limitations. First, it does not consider indirect interactions. Indirect interactions in this food web likely impact fitness outcomes through both density and trait mediated effects. Second, our study does not explicitly incorporate environmental gradients. Comparing across environmental gradients can provide more nuanced insights into context-dependency of animal interactions (McCary & Schmitz 2021). The contradictory patterns of the zoogeochemical impacts of mussels in the microbial part of the aquatic food webs - proliferation of toxic cyanobacterial blooms in Lake Erie (Vanderploeg *et al*. 2009) and the collapse of the spring phytoplankton bloom in Lake Michigan (Vanderploeg *et al*. 2010) - underscores the complex context-dependent nature of this animal-microbe system. We suggest that future research should further investigate the metabolic niche space of these microbial communities, using laboratory-based experiments and population equation models, to explore the underlying eco-evolutionary processes driving diversity patterns.

## Conclusion

Our findings demonstrate that microbial resistance to predation is not encoded by a single gene or metabolic module but rather emerges as a hierarchically structured trait shaped by the interplay between genomic architecture, functional composition, and environmental variation. At the granular protein level (pfam), resistant populations demonstrated independent acquisition of diverse resistance mechanisms. However, functional niche overlap across complex metabolic networks (KEGG) and protein evolution (COG), suggests shared metabolic pathways underpinning resistance at higher levels of functional organization, even across ecosystems with differing nutrient regimes (Denef *et al*. 2016). Moreover, we find no evidence for the existence of a growth-defense tradeoff, likely due to heterogeneous environmental conditions in Lake Erie relaxing equilibrium-based constraints of the growth-defense trade-off theory. Finally, our statistical modelling results, combined with evidence from research in non-equilibrium ecological theory, provide a conceptual framework to explain observed patterns of genetic and functional diversity associated with resistance to mussel predation. Perhaps the following niche-based metaphor best captures the microbial diversity patterns in this predator-prey system: nitrogen and energy metabolism represent key features defining the geometric boundaries of the functional space, environmental gradients carve the adaptive landscape, and mobilome activity - whether through HGT or within genome rearrangements - links diverse ecological strategies across bacterial populations.

### Materials and Methods

We conducted three mussel feeding experiments (June, August, September) in 2019 using water from the western basin of Lake Erie. Our previous work demonstrates that mussel predation altered the structure and composition of bacterial communities in these experiments (Dahal *et al*. 2023). Here, we examine how these predation-induced shifts translate to shifts in bacterial functional traits. Each experiment contained control and mussel treatments, where mussel siphon activity was monitored to confirm feeding. At the end of feeding experiments, water samples were filtered, and DNA was extracted, and libraries were sequenced using Illumina NovaSeq.

Metagenomics reads were processed using the Deneflab metagenomic pipeline: https://computational-boot-camp.readthedocs.io, functional annotation and genome reconstruction was performed using Anvi’o (Eren *et al*. 2015). Bacterial growth rates were estimated using gRodon (intrinsic) and irep (instantaneous) (Brown *et al*. 2016; Weissman *et al*. 2021). Coverage, normalized to the read depth, was used to estimate the relative abundance of metagenome assembled genomes, and classify them as resistant or susceptible populations to mussel predation. Differences in abundance, diversity, and functional traits were assessed using Wilcoxon tests, generalized linear models, and mixed-effects models. Full details of sampling, experimental design, sequencing, bioinformatic workflows, and statistical modelling are provided in the Supplemental Information Methods.

## Supporting information

This file contains detailed methods and supplemental figures and tables.

## Acknowledgments

We thank our collaborators at the Ecosystem Dynamics Branch at NOAA in Ann Arbor, especially Dr. Henry A. Vanderploeg, whose guidance was instrumental in the design and coordination of the experimental setup. We are also grateful to the entire team at GLERL, from the boat crew to the research staff, for their support throughout the study. Special thanks to Saujanya Acharya for assistance with statistical modeling, and to Dr. Jackie Lee Weissman for their support with gRodon. This work was supported by the Michigan Sea Grant Graduate Student Fellowship (2022–2023) and the NOAA Great Lakes Omics program distributed through the UM Cooperative Institute for Great Lakes Research (NA17OAR4320152 and NA22OAR4320150 to V.J.D). This is CIGLR contribution number XXX.

## Notes

**Competing Interest Statement**: The authors declare no conflict of interest.

https://www.ncbi.nlm.nih.gov/bioproject?term=PRJNA1345417&cmd=DetailsSearch

https://github.com/nikeshd/Lake-Erie-Growth-Defense-Tradeoff

### Competing Interest Statement

The authors have declared no competing interest.

## Reference

Acman, M., Wang, R., Van Dorp, L., Shaw, L.P., Wang, Q., Luhmann, N., et al. (2022). Role of mobile genetic elements in the global dissemination of the carbapenem resistance gene blaNDM. Nat Commun, 13, 1131.

Adler, P.B., Salguero-Gómez, R., Compagnoni, A., Hsu, J.S., Ray-Mukherjee, J., Mbeau-Ache, C., et al. (2014). Functional traits explain variation in plant life history strategies. Proc. Natl. Acad. Sci. U.S.A., 111, 740–745.

Allesina, S. & Levine, J.M. (2011). A competitive network theory of species diversity. Proceedings of the National Academy of Sciences, 108, 5638–5642.

Allison, S.D. (2012). A trait-based approach for modelling microbial litter decomposition. Ecology Letters, 15, 1058–1070.

Arnott, D.L. & Vanni, M.J. (1996). Nitrogen and phosphorus recycling by the zebra mussel (Dreissena polymorpha) in the western basin of Lake Erie, 53.

Berkman, P.A., Haltuch, M.A. & Tichich, E. (1998). Zebra mussels invade Lake Erie muds. Nature, 393, 28–29.

Berry, M.A., Davis, T.W., Cory, R.M., Duhaime, M.B., Johengen, T.H., Kling, G.W., et al. (2017). Cyanobacterial harmful algal blooms are a biological disturbance to Western Lake Erie bacterial communities. Environmental Microbiology, 19, 1149–1162.

Botelho, J., Cazares, A. & Schulenburg, H. (2023). The ESKAPE mobilome contributes to the spread of antimicrobial resistance and CRISPR-mediated conflict between mobile genetic elements. Nucleic Acids Research, 51, 236–252.

Braeckman, U., Provoost, P., Gribsholt, B., Van Gansbeke, D., Middelburg, J., Soetaert, K., et al. (2010). Role of macrofauna functional traits and density in biogeochemical fluxes and bioturbation. Mar. Ecol. Prog. Ser., 399, 173–186.

Brown, C.T., Olm, M.R., Thomas, B.C. & Banfield, J.F. (2016). Measurement of bacterial replication rates in microbial communities. Nat Biotechnol, 34, 1256–1263.

Carrick, H.J., VanCuren, C., Slate, A., Denef, V.J., Dahal, N., Carter, G., et al. (2023). Preferential dreissenid mussel grazing on small plankton can favor cyanobacteria: Experimental evidence from western Lake Erie. Aquatic Ecosystem Health & Management, 26, 100–110.

Carter, G.S., Godwin, C.M., Johengen, T.J., Vanderploeg, H.A., Elgin, A.K., Glyshaw, P.W., et al. (2023). Impacts of dreissenid mussel growth and activity on phytoplankton and nutrients in Lake Erie’s western basin. Aquatic Ecosystem Health & Management, 26, 87–99.

Crowther, T.W., Stanton, D.W.G., Thomas, S.M., A’Bear, A.D., Hiscox, J., Jones, T.H., et al. (2013). Top-down control of soil fungal community composition by a globally distributed keystone consumer. Ecology, 94, 2518–2528.

Dahal, N., Glyshaw, P., Carter, G., Vanderploeg, H.A. & Denef, V.J. (2023). Impacts of an invasive filter-feeder on bacterial biodiversity are context dependent. FEMS Microbiology Ecology, 99, fiac149.

Denef, V.J., Carrick, H.J., Cavaletto, J., Chiang, E., Johengen, T.H. & Vanderploeg, H.A. (2017). Lake Bacterial Assemblage Composition Is Sensitive to Biological Disturbance Caused by an Invasive Filter Feeder. mSphere, 2, e00189–17.

Denef, V.J., Kalnejais, L.H., Mueller, R.S., Wilmes, P., Baker, B.J., Thomas, B.C., et al. (2010). Proteogenomic basis for ecological divergence of closely related bacteria in natural acidophilic microbial communities. Proc. Natl. Acad. Sci. U.S.A., 107, 2383–2390.

Denef, V.J., Mueller, R.S., Chiang, E., Liebig, J.R. & Vanderploeg, H.A. (2016). Chloroflexi CL500-11 Populations That Predominate Deep-Lake Hypolimnion Bacterioplankton Rely on Nitrogen-Rich Dissolved Organic Matter Metabolism and C1 Compound Oxidation. Appl Environ Microbiol, 82, 1423–1432.

Eren, A.M., Esen, Ö.C., Quince, C., Vineis, J.H., Morrison, H.G., Sogin, M.L., et al. (2015). Anvi’o: an advanced analysis and visualization platform for ‘omics data. PeerJ, 3, e1319.

Estes, J.A., Terborgh, J., Brashares, J.S., Power, M.E., Berger, J., Bond, W.J., et al. (2011). Trophic Downgrading of Planet Earth. Science, 333, 301–306.

Falkowski, P.G., Fenchel, T. & Delong, E.F. (2008). The Microbial Engines That Drive Earth’s Biogeochemical Cycles. Science, 320, 1034–1039.

Gould, A.L., Zhang, V., Lamberti, L., Jones, E.W., Obadia, B., Korasidis, N., et al. (2018). Microbiome interactions shape host fitness. Proc. Natl. Acad. Sci. U.S.A., 115.

Gravel, D., Albouy, C. & Thuiller, W. (2016). The meaning of functional trait composition of food webs for ecosystem functioning. Phil. Trans. R. Soc. B, 371, 20150268.

Grime, J.P. (1977). Evidence for the Existence of Three Primary Strategies in Plants and Its Relevance to Ecological and Evolutionary Theory. The American Naturalist, 111, 1169–1194.

Hairston, N.G., Smith, F.E. & Slobodkin, L.B. (1960). Community Structure, Population Control, and Competition. The American Naturalist, 94, 421–425.

Havel, J.E., Kovalenko, K.E., Thomaz, S.M., Amalfitano, S. & Kats, L.B. (2015). Aquatic invasive species: challenges for the future. Hydrobiologia, 750, 147–170.

Higgins, S.N. & Vander Zanden, M.J. (2010). What a difference a species makes: a meta–analysis of dreissenid mussel impacts on freshwater ecosystems. Ecological Monographs, 80, 179–196.

Huisman, J., Codd, G.A., Paerl, H.W., Ibelings, B.W., Verspagen, J.M.H. & Visser, P.M. (2018). Cyanobacterial blooms. Nat Rev Microbiol, 16, 471–483.

Hutchinson, G.E. (1961). The Paradox of the Plankton. The American Naturalist, 95, 137–145.

Kanehisa, M. (2019). Toward understanding the origin and evolution of cellular organisms. Protein Science, 28, 1947–1951.

Knutie, S.A., Wilkinson, C.L., Kohl, K.D. & Rohr, J.R. (2017). Early-life disruption of amphibian microbiota decreases later-life resistance to parasites. Nat Commun, 8, 86.

Koonin, E.V. (2005). Orthologs, Paralogs, and Evolutionary Genomics. Annu. Rev. Genet., 39, 309–338.

Lajoie, G. & Kembel, S.W. (2019). Making the Most of Trait-Based Approaches for Microbial Ecology. Trends in Microbiology, 27, 814–823.

Lee, P.O., McLellan, S.L., Graham, L.E. & Young, E.B. (2015). Invasive dreissenid mussels and benthic algae in Lake Michigan: characterizing effects on sediment bacterial communities. FEMS Microbiology Ecology, 91, 1–12.

Lennon, J.T. & Denef, V.J. (2015). Evolutionary Ecology of Microorganisms: From the Tamed to the Wild. In: Manual of Environmental Microbiology (eds. Yates, M.V., Nakatsu, C.H., Miller, R.V. & Pillai, S.D.). ASM Press, Washington, DC, USA, p. 4.1.2-1-4.1.2-12.

Levins, R. (1979). Coexistence in a Variable Environment. The American Naturalist, 114, 765–783.

Li, J., Ianaiev, V., Huff, A., Zalusky, J., Ozersky, T. & Katsev, S. (2021). Benthic invaders control the phosphorus cycle in the world’s largest freshwater ecosystem. Proc. Natl. Acad. Sci. U.S.A., 118, e2008223118.

Litchman, E. & Klausmeier, C.A. (2008). Trait-Based Community Ecology of Phytoplankton. Annu. Rev. Ecol. Evol. Syst., 39, 615–639.

Malik, A.A., Martiny, J.B.H., Brodie, E.L., Martiny, A.C., Treseder, K.K. & Allison, S.D. (2020). Defining trait-based microbial strategies with consequences for soil carbon cycling under climate change. The ISME Journal, 14, 1–9.

Martiny, A.C., Treseder, K. & Pusch, G. (2013). Phylogenetic conservatism of functional traits in microorganisms. The ISME Journal, 7, 830–838.

Mazorra-Alonso, M., Tomás, G. & Soler, J.J. (2021). Microbially Mediated Chemical Ecology of Animals: A Review of Its Role in Conspecific Communication, Parasitism and Predation. Biology, 10, 274.

McCary, M.A. & Schmitz, O.J. (2021). Invertebrate functional traits and terrestrial nutrient cycling: Insights from a global meta-analysis. Journal of Animal Ecology, 90, 1714–1726.

McFall-Ngai, M., Hadfield, M.G., Bosch, T.C.G., Carey, H.V., Domazet-Lošo, T., Douglas, A.E., et al. (2013). Animals in a bacterial world, a new imperative for the life sciences. Proc. Natl. Acad. Sci. U.S.A., 110, 3229–3236.

McGill, B., Enquist, B., Weiher, E. & Westoby, M. (2006). Rebuilding community ecology from functional traits. Trends in Ecology & Evolution, 21, 178–185.

Metcalf, C.J.E., Henry, L.P., Rebolleda-Gómez, M. & Koskella, B. (2019). Why Evolve Reliance on the Microbiome for Timing of Ontogeny? mBio, 10, 10.1128/mbio.01496-19.

Oksanen, L., Fretwell, S.D., Arruda, J. & Niemela, P. (1981). Exploitation Ecosystems in Gradients of Primary Productivity. The American Naturalist, 118, 240–261.

Pacciani-Mori, L., Giometto, A., Suweis, S. & Maritan, A. (2020). Dynamic metabolic adaptation can promote species coexistence in competitive microbial communities. PLoS Comput Biol, 16, e1007896.

Paine, R.T. (1966). Food Web Complexity and Species Diversity. The American Naturalist, 100, 65–75.

Polz, M.F., Alm, E.J. & Hanage, W.P. (2013). Horizontal gene transfer and the evolution of bacterial and archaeal population structure. Trends in Genetics, 29, 170–175.

Ramin, K.I. & Allison, S.D. (2019). Bacterial Tradeoffs in Growth Rate and Extracellular Enzymes. Front. Microbiol., 10, 2956.

Rodriguez-Valera, F., Martín-Cuadrado, A.-B., Rodriguez-Brito, B., Pasic, L., Thingstad, T.F., Rohwer, F., et al. (2009). Explaining microbial population genomics through phage predation. Nat Prec.

Rohlfs, M. & Churchill, A.C.L. (2011). Fungal secondary metabolites as modulators of interactions with insects and other arthropods. Fungal Genetics and Biology, 48, 23–34.

Scales, N.C., Chase, A.B., Finks, S.S., Malik, A.A., Weihe, C., Allison, S.D., et al. (2022). Differential Response of Bacterial Microdiversity to Simulated Global Change. Appl Environ Microbiol, 88, e02429–21.

Scheffer, M., Rinaldi, S., Huisman, J. & Weissing, F.J. (2003). Why plankton communities have no equilibrium: solutions to the paradox. Hydrobiologia, 491, 9–18.

Schmitz, O.J. & Leroux, S.J. (2020). Food Webs and Ecosystems: Linking Species Interactions to the Carbon Cycle. Annu. Rev. Ecol. Evol. Syst., 51, 271–295.

Schmitz, O.J., Wilmers, C.C., Leroux, S.J., Doughty, C.E., Atwood, T.B., Galetti, M., et al. (2018). Animals and the zoogeochemistry of the carbon cycle. Science, 362, eaar3213.

Song, J., Tang, Z., Zhao, X., Yin, Y., Li, X., Chen, F., et al. (2023). Red imported fire ant nesting affects the structure of soil microbial community. Front. Cell. Infect. Microbiol., 13, 1221996.

Soucy, S.M., Huang, J. & Gogarten, J.P. (2015). Horizontal gene transfer: building the web of life. Nat Rev Genet, 16, 472–482.

Stearns, S.C. (1989). Trade-Offs in Life-History Evolution. Functional Ecology, 3, 259.

Strayer, D.L. (2009). Twenty years of zebra mussels: lessons from the mollusk that made headlines. Frontiers in Ecol & Environ, 7, 135–141.

Szappanos, B., Fritzemeier, J., Csörgő, B., Lázár, V., Lu, X., Fekete, G., et al. (2016). Adaptive evolution of complex innovations through stepwise metabolic niche expansion. Nat Commun, 7, 11607.

Tang, H., Vanderploeg, H.A., Johengen, T.H. & Liebig, J.R. (2014). Quagga mussel (Dreissena rostriformis bugensis) selective feeding of phytoplankton in Saginaw Bay. Journal of Great Lakes Research, 40, 83–94.

Tilman, D. (2000). Causes, consequences and ethics of biodiversity. Nature, 405, 208–211.

Tobias-Hünefeldt, S.P., Wenley, J., Baltar, F. & Morales, S.E. (2021). Ecological drivers switch from bottom–up to top–down during model microbial community successions. The ISME Journal, 15, 1085–1097.

Vandermeer, J. & Perfecto, I. (2023). Intransitivity as a dynamic assembly engine of competitive communities. Proc. Natl. Acad. Sci. U.S.A., 120, e2217372120.

Vandermeer, J. & Perfecto, I. (2024). Combining intransitive and higher-order effects in a coupled oscillator framework: A case study of an ant community. Ecology, 105, e4218.

Vanderploeg, H.A., Glyshaw, P.W., Carrick, H.J., Carter, G.S., Dahal, N., Denef, V.J., et al. (2023). Seasonal interactions between Quagga Mussel grazing and phytoplankton in western Lake Erie: The view from different measuring technologies. Aquatic Ecosystem Health & Management, 26, 111–119.

Vanderploeg, H.A., Johengen, T.H. & Liebig, J.R. (2009). Feedback between zebra mussel selective feeding and algal composition affects mussel condition: did the regime changer pay a price for its success? Freshwater Biology, 54, 47–63.

Vanderploeg, H.A., Liebig, J.R., Carmichael, W.W., Agy, M.A., Johengen, T.H., Fahnenstiel, G.L., et al. (2001). Zebra mussel ( Dreissena polymorpha ) selective filtration promoted toxic Microcystis blooms in Saginaw Bay (Lake Huron) and Lake Erie. Can. J. Fish. Aquat. Sci., 58, 1208–1221.

Vanderploeg, H.A., Liebig, J.R., Nalepa, T.F., Fahnenstiel, G.L. & Pothoven, S.A. (2010). Dreissena and the disappearance of the spring phytoplankton bloom in Lake Michigan. Journal of Great Lakes Research, The Lower Food Web of Lake Michigan: Long-term Trends and the Dreissenid Impact, 36, 50–59.

Vanni, M.J. (2002). Nutrient Cycling by Animals in Freshwater Ecosystems. Annual Review of Ecology, Evolution, and Systematics, 33, 341–370.

Vanni, M.J. (2021). Invasive mussels regulate nutrient cycling in the largest freshwater ecosystem on Earth. Proc. Natl. Acad. Sci. U.S.A., 118, e2100275118.

Veseli, I., Chen, Y.T., Schechter, M.S., Vanni, C., Fogarty, E.C., Watson, A.R., et al. (2024). Microbes with higher metabolic independence are enriched in human gut microbiomes under stress.

Walters, A.W., Hughes, R.C., Call, T.B., Walker, C.J., Wilcox, H., Petersen, S.C., et al. (2020). The microbiota influences the Drosophila melanogaster life history strategy. Molecular Ecology, 29, 639–653.

Watson, A.R., Füssel, J., Veseli, I., DeLongchamp, J.Z., Silva, M., Trigodet, F., et al. (2023). Metabolic independence drives gut microbial colonization and resilience in health and disease. Genome Biol, 24, 78.

Weissman, J.L., Hou, S. & Fuhrman, J.A. (2021). Estimating maximal microbial growth rates from cultures, metagenomes, and single cells via codon usage patterns. Proc. Natl. Acad. Sci. U.S.A., 118, e2016810118.

Werner, E.E. (1986). Amphibian Metamorphosis: Growth Rate, Predation Risk, and the Optimal Size at Transformation. The American Naturalist, 128, 319–341.

Wiedenbeck, J. & Cohan, F.M. (2011). Origins of bacterial diversity through horizontal genetic transfer and adaptation to new ecological niches. FEMS Microbiol Rev, 35, 957–976.

Wyness, A.J., Fortune, I., Blight, A.J., Browne, P., Hartley, M., Holden, M., et al. (2021). Ecosystem engineers drive differing microbial community composition in intertidal estuarine sediments. PLoS ONE, 16, e0240952.

Xia, L., Miao, Y., Cao, A., Liu, Y., Liu, Z., Sun, X., et al. (2022). Biosynthetic gene cluster profiling predicts the positive association between antagonism and phylogeny in Bacillus. Nat Commun, 13, 1023.

