## Supplementary material for "Resistance to Predation is Not Constrained by Growth-Defense Trade-Offs in Freshwater Bacterial Communities": This file contains detailed methods and supplemental figures and tables.

**This PDF file includes:**

Supporting text
Figures S1 to S3
Tables S1 to S4
SI References

#### Supporting Information Text

##### Methods

In this study, we report metagenomic data from three mussel feeding experiments (July 18, August 13, and September 18, 2019) conducted using water collected from the western basin of Lake Erie.

###### *1. Sample collection and experimental design*

Approximately 450 L of water and mussels were collected from the WE4 station (41° 49' 35.4" N and 83° 11' 47.4" W), a long-term monitoring site maintained by NOAA's Great Lakes Environmental Research Laboratory, one day prior to the feeding experiments. Mussels were cleaned of shell debris and kept in lake water overnight in 90-L drums to allow for acclimation. Before the experiments, mussels were transferred to a 50-L aquarium containing lake water for 2 hours to allow acclimatization to the food source, and then sorted into mesh baskets for easy transfer during the experiments (1). We conducted feeding experiments in 20-L high-density polyethylene (HDPE) containers, which were cleaned with bleach, rinsed with MilliQ water, and dried prior to use. Each experiment included four treatments (20–25 quagga mussels per container), while control treatments (containers without mussels) varied across dates: three controls in July and September, and four controls in August. Due to the higher chlorophyll concentration in August, we included two sample types each: whole lake water and diluted lake water (whole water + 0.2 μm filtered lake water). Dilution adjusted final chlorophyll-a (chl-a) concentrations to ~4 μg/L, below the incipient limiting concentration, to ensure feeding effects were detectable and not masked by high chl-a levels. After quagga mussels were added to the treatment containers, their siphon opening time (5–15 minutes after addition) was visually recorded as an indicator of active feeding and experiments began when all treatments indicated active feeding. Chl-a depletion was measured hourly using a bbe-Moldaenke FluoroProbe II fluorometer, with experiments running for 2 hours in July and September and 3 hours in August, determined by the observed feeding rates of mussels.

###### *2. DNA extraction and shotgun sequencing*

At the end of each experiment, we sampled 2 L of water from each treatment using sterile Nalgene bottles. Bacterial communities in these samples were collected onto 47-mm, 0.22-μm PES Millipore Express filters using portable Masterflex peristaltic pumps with Pall polycarbonate filter holders. Filtration was conducted until 2 L of water was processed or for a maximum time of 10 minutes. Filters containing the bacterial samples were then transferred into sterile cryovials, immediately frozen in liquid nitrogen, and stored at -80°C. In the laboratory, DNA was extracted from the filters using the DNeasy Blood & Tissue kit with a modified Qiagen AllPrep Universal kit protocol (2). The extracted DNA was sent to the Advanced Genomics Core at the University of Michigan for shotgun sequencing. At the core, DNA quality was assessed using the Qubit dsDNA High Sensitivity Assay (Fisher Scientific) and TapeStation Genomic DNA reagents (Agilent, Santa Clara, CA). Libraries were prepared with the NEBNext Ultra II FS DNA Library Prep Kit for Illumina (#E7805L, New England Biolabs) following the manufacturer's protocol, using 100–200 ng of DNA input and size-selecting for a 275–745 bp insert size. Library quality was verified using the Qubit dsDNA High Sensitivity Assay (Fisher), LabChip (Perkin Elmer), and a MiSeq run to confirm library balance. Sequencing was performed on an Illumina NovaSeq S4 platform in a 300-cycle run (150 paired-end), following the manufacturer's protocol. Demultiplexed Fastq files were generated using Bcl2fastq2 Conversion Software (Illumina) for downstream analyses.

###### *3. Bioinformatics*

*3.1 Metagenomic assembly and binning*: Data processing was performed using the Deneflab metagenomic pipeline designed for a reproducible and modular workflow: https://computational-boot-camp.readthedocs.io. In brief, bbtools was used for quality control to trim adapters, remove poor-quality reads, and identify optical duplicates (https://sourceforge.net/projects/bbmap). One-third of quality-controlled reads were randomly subsampled, pooled together within the treatment type (control vs. quagga), and assembled using MEGAHIT to create two independent assemblies for each date. Contiguous sequences longer than 2.5 kbp were then used to map reads to the reference assemblies using bwa (3). CONCOCT was used for automated binning (4) and the quality of the bins was assessed using CheckM (5). To optimize mapping and binning performance, the quality of bins was compared using two approaches: (a) mapping reads from 2 samples (1 control and 1 quagga for each date) to the corresponding assembly, and (b) mapping reads from all samples (across dates) to the assembly. We observed that using reads from all samples produced high quality bins and thus, adopted this approach. Then, bins with low redundancy (<10%) and high completion (>80%) were manually refined using Anvi'o (6). Refinement history for each bin is available in the supplementary materials.

*3.2 Dereplication and functional annotation*: Manually curated bins were renamed as metagenome-assembled genomes (MAGs), assigned unique identifiers, grouped into a collection, and exported using the *anvi-summarize* function to generate fasta files and other metadata. To avoid treating highly similar MAGs as distinct populations, dereplication was performed across assemblies using the workflow and scripts from Evans & Denef (2020). Sourmash was used for hierarchical clustering and a multistep pyANI process was run to compute average nucleotide identity (ANI) between MAGs. Dereplication was tested across three coverage thresholds (10%, 25%, and 50%) while maintaining ANI threshold of 99%. The 50% coverage threshold was selected as it provided the balance between reducing redundancy and preserving MAG quality. Three high-quality MAGs that did not pass the 50% threshold were kept due to their high completeness and coverage similarity, potentially representing closely related populations or important genomic diversity patterns. The dereplicated MAGs were then concatenated into a single fasta file for subsequent competitive mapping of quality controlled reads using bwa to quantify coverage values for each population. Competitively mapped coverage of the dereplicated MAGs was used to create a new Anvi'o database following instructions at https://merenlab.org/2016/06/22/anvio-tutorial-v2/. Gene annotation was performed using the following Anvi'o functions: a) *anvi-run-hmms* for Hidden Markov Model (HMM) hits (b) *anvi-run-scg-taxonomy* to affiliate single-copy core genes with taxonomy (c) *anvi-run-ncbi-cogs* for annotation based on NCBI's clusters of orthologous groups, and (e) *anvi-run-kegg-kofams* for annotation based on KEGG Orthologs (KOs). Functional inferences were made using *anvi-run-pfams* to associate genes with the protein family database (European Bioinformatics Institute) and *anvi-estimate-metabolism* to reconstruct metabolic pathways and estimate their completeness within each MAG. The *anvi-summarize function* was used to generate summaries of mean coverage, GC content, genome length, detection, and other metrics. Biosynthetic gene clusters (BGCs) in resistant MAGs were characterized using antiSMASH with default parameters (8).

*3.3 Phylogenetic tree:* A phylogenetic tree was constructed using six ribosomal proteins (Ribosomal_L1, L2, L3, L4, L5, L6), in alignment with standard practices in microbial phylogenomics (9). These genes were concatenated and phylogenetic distances were computed in Anvi'o using the *anvi-gen-phylogenomic-tree* function, which uses FastTree. Newick tree was then visualized using iTOL.

*3.4 Growth Rate Estimation:* Maximum growth rates were predicted for all resistant and susceptible populations (MAGs) using gRodon, an R package that estimates growth rates based on the distance between features in the genome-wide codon usage bias space (10). Instantaneous growth rates were estimated using iRep, which leverages mean coverage differences between the origin of replication and the terminus of bacterial populations to estimate replication index (11). A total of 574 irep runs were performed on 30 populations (15 resistant and 15 randomly selected susceptible) to ensure irep value for a population was estimated for each treatment and date whenever the population was present (across a total of 22 assemblies). For any given date, at least 22 out of 30 populations were present.

##### *4. Statistical analyses and modelling*

*4.1 Abundance quantification*: To determine a presence threshold for bacterial populations, we plotted detection vs. coverage for all samples. Detection increased with coverage but started to asymptote at ~70%. Thus, populations with detection below 0.70 were excluded to minimize stochastic shifts at low abundance levels. Relative abundance was quantified using mean coverage q2q3 values estimated by Anvi'o. The q2q3 values represent the mean genome coverage (total coverage of all nucleotides divided by the contig length), excluding outliers in the first and third quartiles. These values were normalized by sequencing depth across samples to ensure consistent relative abundance estimates. When analyzing resistance as a continuous variable, the ratio of mean relative abundance between quagga mussel and control treatments (Q:C ratio) was used. For categorical comparisons, populations were classified as resistant or susceptible based on the consistency of their response to mussel predation across dates: (1) a population was classified as resistant if its mean relative abundance was consistently higher in the mussel treatment compared to control whenever it showed a significant response and (2) a population was classified as susceptible if its mean relative abundance was significant and consistently lower in the mussel treatment. Statistical significance was measured using a generalized linear model (GLM, negative binomial family) and only populations with consistent response were retained for categorical comparisons.

*4.2 Comparative trait analyses:* We assessed differences between resistant and susceptible populations across multiple genomic and functional traits: *(a) Genomic traits:* differences in the median GC content and genome length were tested using phylogenetic least squares regression (PGLS) and Wilcoxon tests. *(b) Maximum growth rates:* predicted from gRodon were compared using the Wilcoxon test. *(c) Protein family (pfam) profiles:* mean relative abundance of pfams shared between resistant and susceptible populations were compared using Wilcoxon tests, followed by Benjamini-Hochberg corrections to account for false discovery rates *(d) Metabolic module profiles:* differential abundance of KEGG metabolic modules between the two populations was computed using *anvi-compute-enrichment* function (12). *(e) Metabolic module completeness:* stepwise mean module completeness scores were compared for significantly overrepresented metabolic modules using Wilcoxon tests. *(f) Temporal persistence: relative* proportion of resistant populations across dates was compared using chi-square test.

*4.3 Diversity analyses:* Alpha and beta diversity analyses were performed using the vegan package. Principal Coordinate Analysis (PCoA) was used to visualize differences in bacterial community composition.

*4.4 Statistical modelling of growth rates:* To evaluate correlation between resistance and observed instantaneous growth rates, linear mixed-effects models (LMMs) were implemented using the lmer package. Baseline model (model 1) included population (MAG) as a random intercept, and population category (resistant vs. susceptible) and treatment as fixed effects. Interaction model (model 2) allowed for interaction between population category and treatment, and added date as fixed effect along with population as a random slope. The full model included both treatment and date as random slopes, allowing growth rates to vary flexibly across time. Model selection was performed using ANOVA, and model 2 and model 3 were tested for explanatory power and robustness by training the models on decreasing proportions of data (80, 70, 60, 50, and 40%).

model 1: growth_rate ~ population_category + treatment + (1 | MAG)
model 2: growth_rate ~ population_category X treatment + date + (1 + treatment | MAG)
model 3: growth_rate ~ population_category X treatment + treatment + date + (1+ treatment + date | MAG)

All statistical analyses were performed in RStudio (using vegan, nlme, lmer, MASS, ape, ggtree, phytools, gRodon packages). R scripts are available here: https://github.com/nikeshd/Lake-Erie-Growth-Defense-Tradeoff

### Figures
#### Figure S1

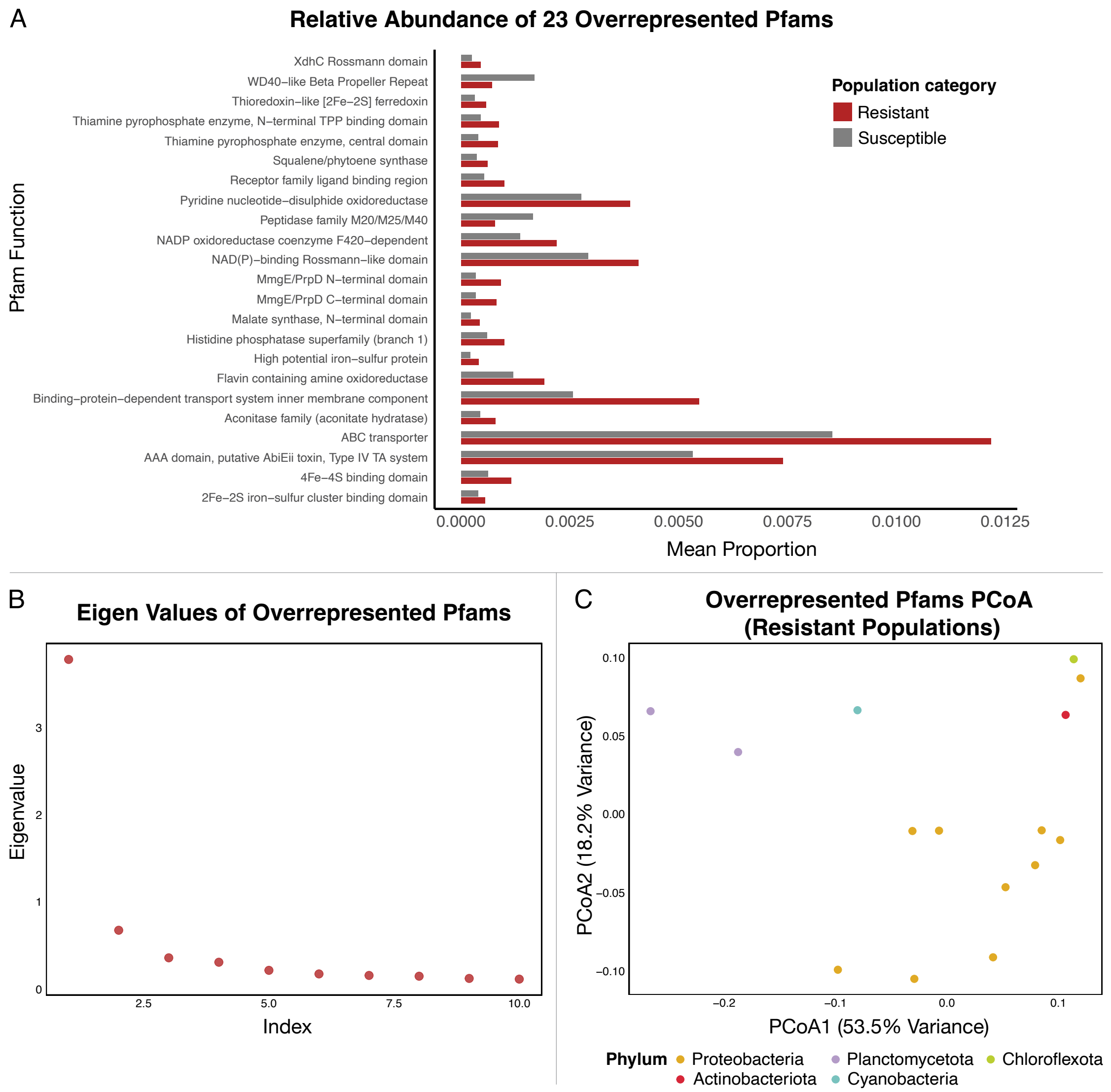


Patterns of variation in pfam composition. (A) Relative abundance of 23 pfams overrepresented in resistant (red) and susceptible (gray) populations. (B) Top 10 eigenvalues from the Principal Coordinates Analysis (PCoA, using Bray-Curtis) of pfam composition for overrepresented pfams (Figure 2C) showing that most variation is captured by the first eigenvalue. (C) PCoA (Bray-Curtis) of 23 pfam composition in resistant populations. There was a significant clustering by the taxonomic structure (PERMANOVA, p-value = 0.002) but significant dispersion differences (Anova, p-value = 0.012) also contributed to the observed clustering pattern.

#### Figure S2

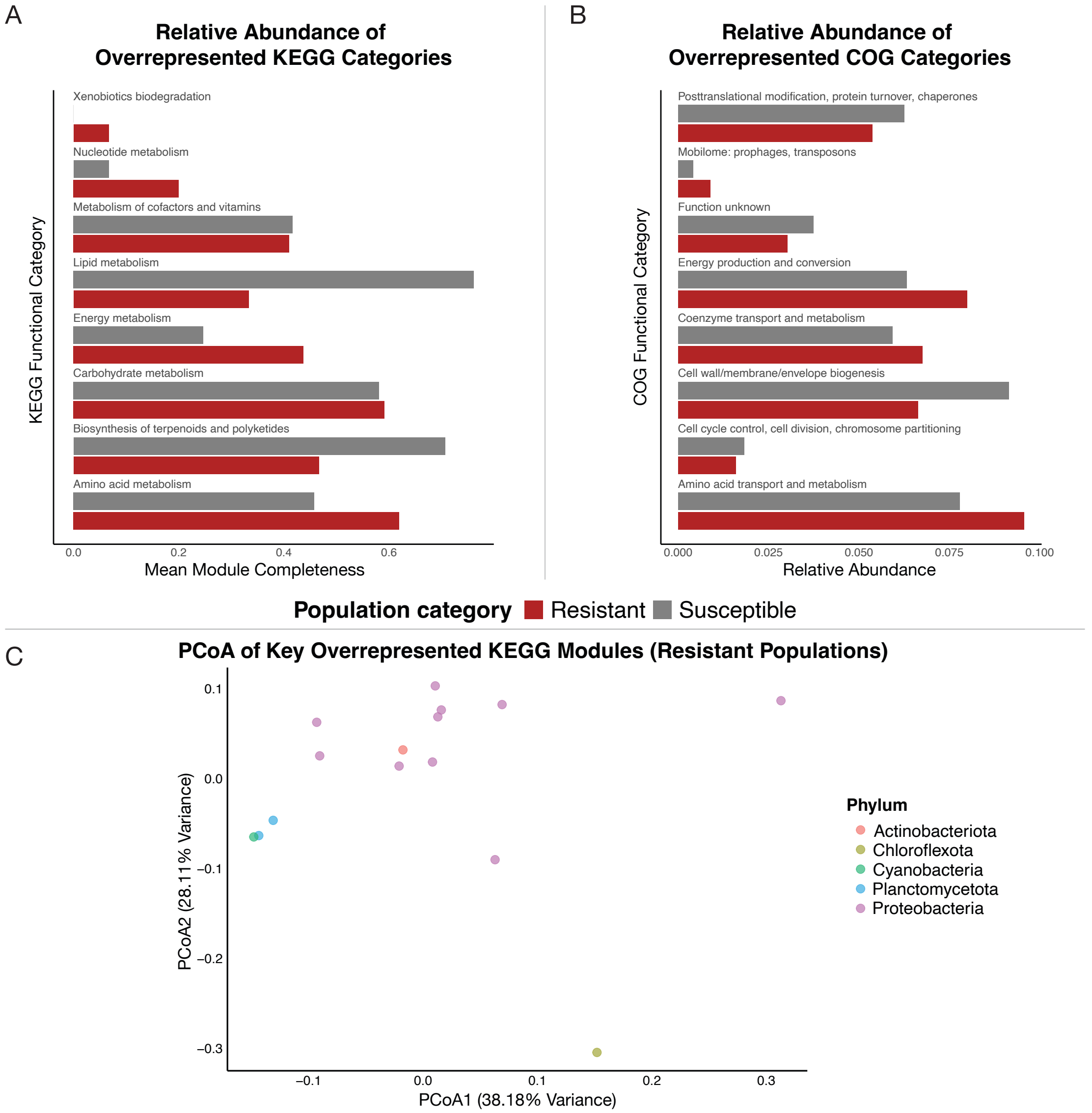


Patterns of variation in the composition of broad functional categories. A) Relative abundance of 35 overrepresented functional categories in resistant (red) and susceptible (gray) populations, grouped by broad metabolic pathways. (B) Relative abundance of 8 overrepresented COG functional categories in the two populations. (C) PCoA (Bray-Curtis) of metabolic module composition for KEGG pathways related to amino acid and energy metabolism in resistant populations. There was a marginally significant clustering by the taxonomic structure (PERMANOVA, p-value = 0.062) but significant dispersion differences (ANOVA, p-value = 0.043) also contributed to the observed clustering pattern.

**Figure S3**

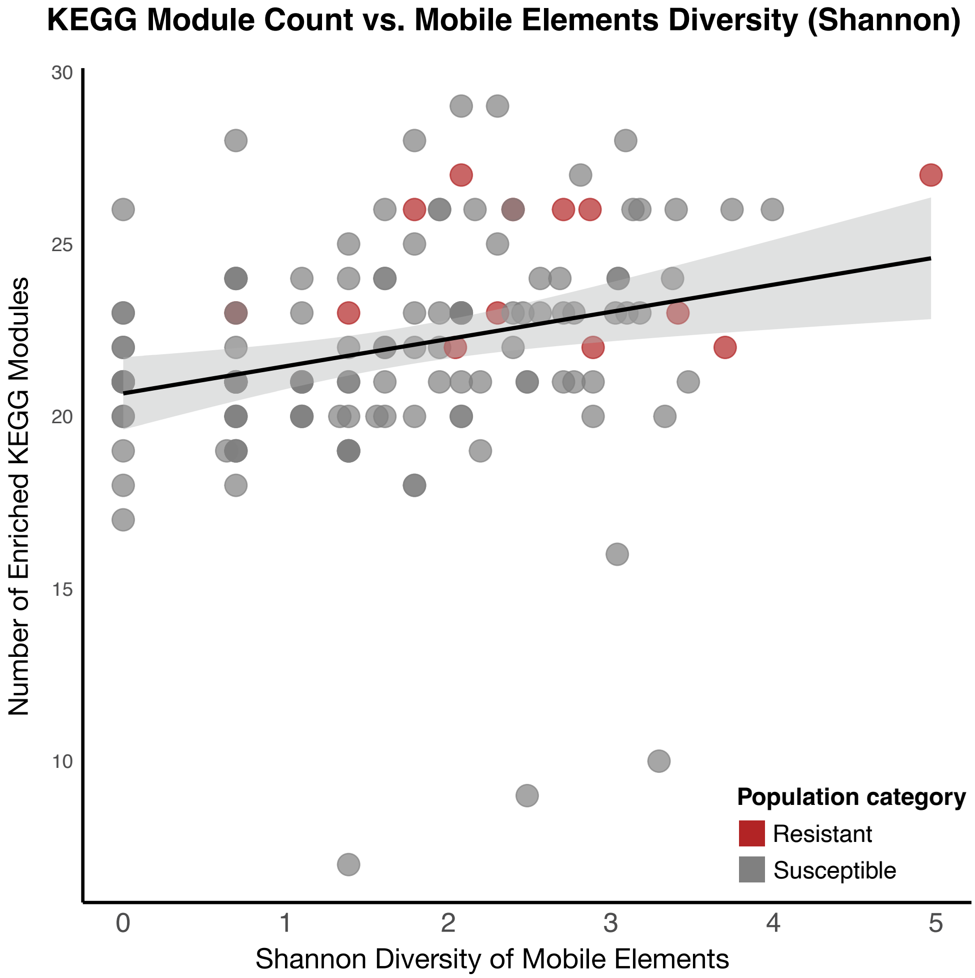


Scatter plot showing the relationship between Shannon diversity of the mobilome COG category (X-axis) and the number of differentially abundant KEGG metabolic pathway modules (Y-axis) for both, resistant (red) and susceptible, populations. A significant positive correlation was observed (Spearman's $\rho = 0.36$, $p < 0.001$), indicating that populations with greater mobilome diversity tend to have a higher number of overrepresented metabolic modules.

**Tables**

**Table S1**. Summary of 181 dereplicated MAGs retained from the Lake Erie bacterial communities across dates and experimental treatments (available here: https://github.com/nikeshd/Lake-Erie-Growth-Defense-Tradeoff/blob/main/R%20output/mag_summary.csv)

**Table S2.** Summary of fixed and random effects from statistical models used to fit instantaneous growth rates estimated from iRep.

| Model | Fixed Effects (Estimate ± SE) | Random Effects (Variance) |
|---|---|---|
| Model 1 | - Intercept: 2.67 ± 0.15***<br>- Population category: -0.24 ± 0.21 (NS)<br>- Treatment: 0.39 ± 0.05*** | - MAG: 0.303<br>- Residual: 0.418 |
| Model 2 | - Intercept: 2.61 ± 0.14***<br>- Population category: -0.51 ± 0.19*<br>- Mussel Treatment: 0.21 ± 0.10*<br>- Date:<br>• Aug: = 0.209 ± 0.067**<br>• Sep: = 0.242 ± 0.069*** | - MAG: 0.219<br>- Residual: 0.379 |
| Model 3 | - Intercept: 2.68 ± 0.13***<br>- Population category: -0.57 ± 0.18**<br>- Mussel Treatment: 0.23 ± 0.10*<br>- Population category × mussel treatment: 0.43 ± 0.15**<br>- Date:<br>• Aug: 0.162 ± 0.152 (NS)<br>• Sep: 0.166 ± 0.171 (NS) | - MAG: 0.260<br>- Treatment: 0.140<br>- Date:<br>• Aug: 0.52<br>• Sep: 0.685<br>- Residual: 0.178 |

**Table S3**
Summary of the performance results comparing model 2 and model 3 for robustness and accuracy.

| Training data (proportion) | Model | Mean Squared Error (MSE) | R2 |
|---|---|---|---|
| 0.8 | model 2 | 0.449 | 0.418 |
| 0.8 | model 3 | 0.237 | 0.692 |
| 0.7 | model 2 | 0.517 | 0.444 |
| 0.7 | model 3 | 0.293 | 0.68 |
| 0.6 | model 2 | 0.582 | 0.36 |
| 0.6 | model 3 | 0.308 | 0.665 |
| 0.5 | model 2 | 0.387 | 0.443 |
| 0.5 | model 3 | 0.232 | 0.667 |
| 0.4 | model 2 | 0.472 | 0.380 |
| 0.4 | model 3 | 0.293 | 0.616 |

**Table S4**
List of the biosynthetic gene clusters (BGCs) identified in resistant populations using anti-SMASH

| MAG_ID | BGC_Type | Start | End | t_phylum | t_genus |
|---|---|---|---|---|---|
| ND35_MAG_00023 | terpene | 0 | 14839 | Proteobacteria | UBA954 |
| ND35_MAG_00042 | terpene | 0 | 10132 | Proteobacteria | CAINMN01 |
| ND42_MAG_00043 | terpene | 3413 | 17124 | Proteobacteria | UBA954 |
| ND42_MAG_00043 | terpene | 0 | 14953 | Proteobacteria | UBA954 |
| ND42_MAG_00067 | terpene | 0 | 15363 | Proteobacteria | Fonsibacter |
| ND50_MAG_00002 | terpene | 116284 | 138285 | Planctomycetota | UBA6894 |
| ND50_MAG_00002 | hglE-KS | 7061 | 60153 | Planctomycetota | UBA6894 |
| ND50_MAG_00002 | terpene | 224866 | 246834 | Planctomycetota | UBA6894 |
| ND50_MAG_00002 | terpene | 81837 | 103670 | Planctomycetota | UBA6894 |
| ND50_MAG_00002 | T3PKS | 6922 | 40188 | Planctomycetota | UBA6894 |
| ND50_MAG_00021 | terpene | 0 | 13118 | Proteobacteria | CAIUPE01 |
| ND50_MAG_00021 | hserlactone | 0 | 12887 | Proteobacteria | CAIUPE01 |
| ND50_MAG_00044 | linaridin | 0 | 14233 | Proteobacteria | CAINMN01 |
| ND50_MAG_00044 | redox-cofactor | 13255 | 26579 | Proteobacteria | CAINMN01 |
| ND50_MAG_00044 | terpene | 0 | 10091 | Proteobacteria | CAINMN01 |
| ND50_MAG_00055 | terpene | 53377 | 74280 | Actinobacteriota | Planktophila |
| ND57_MAG_00023 | terpene | 0 | 17968 | Planctomycetota | Pirellula |
| ND57_MAG_00023 | NRPS-like | 0 | 25513 | Planctomycetota | Pirellula |
| ND57_MAG_00023 | terpene | 0 | 20133 | Planctomycetota | Pirellula |
| ND57_MAG_00023 | T3PKS | 0 | 30344 | Planctomycetota | Pirellula |
| ND57_MAG_00061 | T1PKS, NRPS, NRPS-like | 0 | 67595 | Cyanobacteria | Microcystis |
| ND57_MAG_00061 | T1PKS, NRPS-like, T3PKS | 0 | 26162 | Cyanobacteria | Microcystis |
| ND57_MAG_00061 | terpene | 6373 | 18536 | Cyanobacteria | Microcystis |
| ND57_MAG_00061 | NRPS, NRPS-like | 0 | 13127 | Cyanobacteria | Microcystis |
| ND57_MAG_00061 | LAP | 0 | 14421 | Cyanobacteria | Microcystis |
| ND57_MAG_00061 | T1PKS | 0 | 14025 | Cyanobacteria | Microcystis |

#### SI References:


1. H. Tang, H. A. Vanderploeg, T. H. Johengen, J. R. Liebig, Quagga mussel (Dreissena rostriformis bugensis) selective feeding of phytoplankton in Saginaw Bay. *Journal of Great Lakes Research* **40**, 83–94 (2014).
2. A. McCarthy, E. Chiang, M. L. Schmidt, V. J. Denef, RNA Preservation Agents and Nucleic Acid Extraction Method Bias Perceived Bacterial Community Composition. *PLOS ONE* **10**, e0121659 (2015).
3. H. Li, R. Durbin, Fast and accurate short read alignment with Burrows–Wheeler transform. *Bioinformatics* **25**, 1754–1760 (2009).
4. J. Alneberg, *et al.*, CONCOCT: Clustering cONtigs on COverage and ComposiTion. [Preprint] (2013). Available at: http://arxiv.org/abs/1312.4038 [Accessed 10 February 2025].
5. D. H. Parks, M. Imelfort, C. T. Skennerton, P. Hugenholtz, G. W. Tyson, CheckM: assessing the quality of microbial genomes recovered from isolates, single cells, and metagenomes. *Genome Res.* **25**, 1043–1055 (2015).
6. A. M. Eren, *et al.*, Anvi'o: an advanced analysis and visualization platform for 'omics data. *PeerJ* **3**, e1319 (2015).
7. J. T. Evans, V. J. Denef, To Dereplicate or Not To Dereplicate? *mSphere* **5**, e00971-19 (2020).
8. K. Blin, *et al.*, antiSMASH 6.0: improving cluster detection and comparison capabilities. *Nucleic Acids Research* **49**, W29–W35 (2021).
9. L. A. Hug, *et al.*, A new view of the tree of life. *Nat Microbiol* **1**, 1–6 (2016).
10. J. L. Weissman, S. Hou, J. A. Fuhrman, Estimating maximal microbial growth rates from cultures, metagenomes, and single cells via codon usage patterns. *Proc. Natl. Acad. Sci. U.S.A.* **118**, e2016810118 (2021).
11. C. T. Brown, M. R. Olm, B. C. Thomas, J. F. Banfield, Measurement of bacterial replication rates in microbial communities. *Nat Biotechnol* **34**, 1256–1263 (2016).
12. A. Shaiber, *et al.*, Functional and genetic markers of niche partitioning among enigmatic members of the human oral microbiome. *Genome Biol* **21**, 292 (2020).
